# Structural basis for the folding of PINK1 by the HSP90–CDC37 chaperone complex

**DOI:** 10.1101/2025.10.17.682828

**Authors:** Kei Okatsu, Hayato Yamamoto, Akinori Okamoto, Shinya H. Goto, Yumiko Nishimoto, Yukihiko Sugita, Takeshi Noda, Shuya Fukai

**Affiliations:** Department of Chemistry, Graduate School of Science, Kyoto University, Kyoto, 606-8502, Japan; Laboratory of Ultrastructural Virology, Institute for Life and Medical Sciences, Kyoto University, Kyoto, Japan; Laboratory of Ultrastructural Virology, Graduate School of Biostudies, Kyoto University, Kyoto, Japan

**Author notes:** Correspondence should be addressed to K.O. and S.F.

## Abstract

PTEN-induced kinase 1 (PINK1) is a mitochondrial serine/threonine kinase that plays a central role in Parkin-dependent mitophagy. Mutations in PINK1 are associated with familial Parkinson’s disease. PINK1 is a high-affinity client of the HSP90–CDC37 complex and is stabilized by this chaperone system. However, the molecular mechanism by which HSP90–CDC37 facilitates the folding of PINK1 remains unclear. Here, we present a cryogenic electron microscopy structure of the human PINK1–HSP90–CDC37 complex. The β5 strand of the PINK1 N-lobe is accommodated in the central channel of the HSP90 dimer, which holds the PINK1 kinase domain in a partially unfolded state. The C-lobe and unique C-terminal extension (CTE) of PINK1 is folded. HSP90 covers the CTE of PINK1, which overlaps with interaction sites for TOM5, TOM20, and the PINK1 N-helix. The HPNI motif of CDC37 interacts with the C-lobe of PINK1, mimicking the HPNI motif in the N-lobe. The pathogenic mutation L347P is suggested to disrupt these interactions, while H271Q is located within the HPNI motif in the N-lobe of PINK1. These findings provide structural insights into the folding of PINK1 and its dysfunction in Parkinson’s disease.

## Introduction

Mutations in the genes encoding PINK1 and Parkin (also known as PARK6 and PARK2, respectively) can cause autosomal recessive Parkinson’s disease (1, 2). PINK1 is a mitochondrial serine/threonine kinase that plays a critical role in Parkin-dependent mitophagy to maintain mitochondrial quality (3–5). Under normal conditions, PINK1 is imported into mitochondria in a membrane potential-dependent manner and cleaved by mitochondrial proteases (6–8). During these import and processing steps, the kinase domain of PINK1 remains exposed to the cytosol in an inactive, non-phosphorylated state (9, 10). The cleaved PINK1 is released from mitochondria and subsequently degraded by the ubiquitin–proteasome pathway, resulting in a rapid turnover of PINK1 (3, 4, 11). Proteasome inhibition causes the accumulation of excessive PINK1 in the cytosol and induces the aggregate formation (11, 12). Meanwhile, loss of the membrane potential of damaged mitochondria abrogates the import and processing of PINK1 and facilitates the PINK1 accumulation on the outer mitochondrial membrane to induce Parkin-dependent mitophagy (10). The state of PINK1 is regulated at multiple steps, ensuring precise surveillance of damaged mitochondria.

The accumulated PINK1 on the outer mitochondrial membrane becomes activated through autophosphorylation in trans (10, 13) and phosphorylates both ubiquitin and the ubiquitin-like (UBL) domain of Parkin (14–17). The phosphorylated Parkin interacts with the phosphorylated ubiquitin, which allosterically activates Parkin (16, 18). The activated Parkin promotes polyubiquitination of mitochondrial substrates to initiate the selective clearance of damaged mitochondria (19). A recent cryo-EM study revealed that the PINK1–TOM complex assembles around VDAC2 and promotes activation of PINK1 through trans auto-phosphorylation (20). The C-terminal extension (CTE) of the PINK1 kinase domain, which is unique to PINK1 and characterizes its atypical kinase architecture, interacts with its N-helix as well as with TOM5 and TOM20, which are components of the TOM complex. In addition, the N-helix is also involved in the interaction with TOM20, and mutations in the N-helix such as C125G and Q126P disrupt this interaction (21, 22).

A kinome-wide quantitative analysis of the HSP90–CDC37–client interaction identified PINK1 as a high-affinity substrate of the HSP90–CDC37 complex (23). HSP90 and its co-chaperone CDC37 are a central chaperone complex for protein kinases (24, 25), which is required for the folding and stabilization of a wide range of kinases (23). The stability of PINK1 is influenced by the HSP90–CDC37 chaperone system (26–29). The L347P mutation of PINK1, a Parkinson’s disease–associated mutation, disrupts the PINK1–HSP90–CDC37 interaction and leads to rapid degradation of PINK1 (26, 28). Despite the importance of the PINK1 stabilization by the HSP90–CDC37 chaperone complex, its underlying structural mechanism remains to be elucidated.

In this study, we report the cryo-EM structure of the PINK1–HSP90–CDC37 complex at a nominal resolution of 3.08 Å. The kinase domain of PINK1 is held in an immature conformation. The N-lobe of PINK1 remains unfolded, whereas the C-lobe and CTE of PINK1 are folded. HSP90 covers the CTE of PINK1. The C-lobe of PINK1 interacts with the conserved HPNI motif of CDC37, which mimics the HPNI motif present in the folded kinase domain of PINK1. Pathogenic mutations H271Q and L347P of PINK1 affect intramolecular interactions via the HPNI motif. Furthermore, the L347P mutation in the C-lobe of PINK1 interferes with the intermolecular stabilization mediated by the HPNI motif of CDC37. These findings provide structural insights into the recognition and stabilization of the atypical kinase architecture of PINK1 by the HSP90–CDC37 complex, and highlight structural and mechanistic relationships between chaperone-mediated folding and pathogenic mutations causing Parkinson’s disease.

## Results

### Complex formation between human PINK1 and the HSP90–CDC37 chaperone

PINK1 consists of an N-terminal mitochondrial targeting sequence (MTS), an outer mitochondrial localization signal (OMS), and a single transmembrane domain (TMD) followed by N-helix and a kinase domain (Fig. 1a). The N-lobe of the kinase domain of PINK1 is composed of five β-strands, the αC-helix and three insertions. The C-lobe and the unique CTE is mainly composed of α-helices. The cleft between the N- and C-lobes forms the ATP-binding pocket. The CTE interacts with the N-helix and forms an interface for binding to components of the TOM complex. HSP90 is a dimeric molecular chaperone. Each protomer contains an ATP-binding N-terminal domain (NTD^HSP90^), a middle domain (MD^HSP90^), and a C-terminal dimerization domain (CTD^HSP90^) (Fig. 1a). The NTD^HSP90^ and MD^HSP90^ are connected by a flexible charged linker (FCL^HSP90^). Two protomers form a channel through dimerization. This channel adopts a clamp-like structure and accommodates client proteins (Fig. 1b). CDC37 functions as a kinase-specific co-chaperone. The N-terminal domain of CDC37 (NTD^CDC37^) directly binds to the kinase domain of client proteins (Fig. 1a). The middle domain of CDC37 (MD^CDC37^) connects to HSP90 through a linker region that extends across the outer surface of the HSP90 dimer, thereby bridging kinases to the chaperone (Fig. 1b).

**Figure 1.**
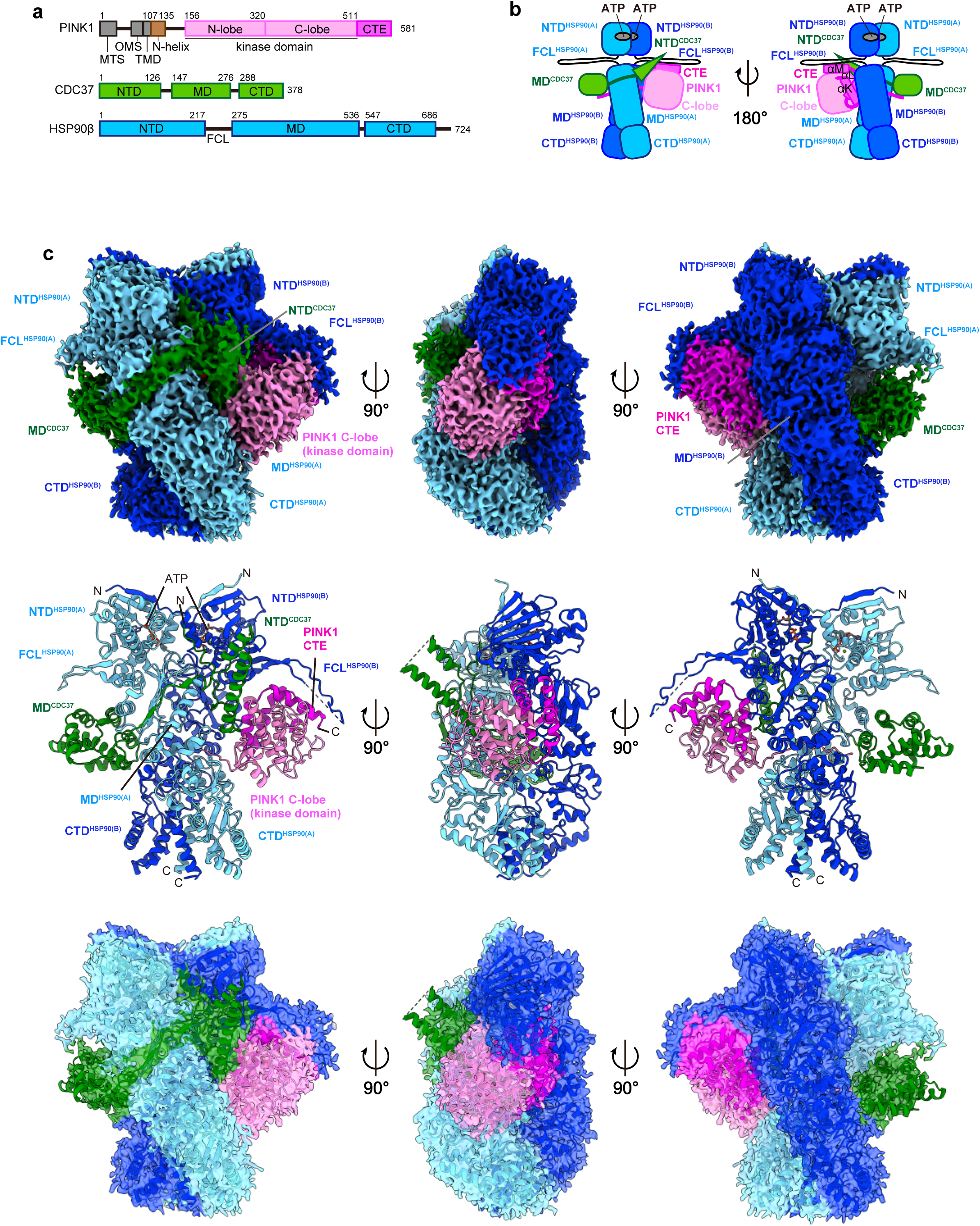
Cryo-EM structure of the PINK1–HSP90–CDC37 complex. **(a)** Domain organizations of human PINK1, CDC37, and HSP90β. MTS, N-terminal mitochondrial targeting sequence; OMS, outer mitochondrial localization signal; TMD, transmembrane domain; CTE, C-terminal extension; NTD, N-terminal domain; MD, middle domain; CTD, C-terminal domain. The PINK1 kinase domain consisting of the N- and C-lobes is colored in pink, the CTE in magenta, CDC37 in green, and HSP90β in cyan. **(b)** Schematic diagram of the PINK1–HSP90–CDC37 complex. The coloring scheme is the same as in (a) except that the two HSP90β protomers are colored in cyan and blue. **(c)** Cryo-EM density map and structure of the PINK1–HSP90–CDC37 complex at 3.08 Å resolution. The coloring scheme is the same as in (b).

PINK1 is cleaved by PARL and exposes Phe104 at the N-terminus (7, 11). The phenylalanine residue at the N-terminus is recognized by the E3 ubiquitin ligases (Ubr1, Ubr2, and Ubr4) of the N-end rule pathway as the destabilizing residue and subsequently degraded by the proteasome (11). To express the stabilized fragment of PINK1, we replaced Phe104 with Met, and deleted the N-terminal 103 residues. PINK1 has been reported to interact with both HSP90 isoforms, HSP90α and HSP90β (27). Because HSP90β is the constitutively expressed isoform and is known to form more stable client–chaperone complexes, we chose HSP90β for this structural study. For cryo-EM analysis, we co-expressed human PINK1 (residues 104–581, F104M), HSP90β, and CDC37 in insect cells and purified the PINK1–HSP90β–CDC37 complex. HSP90 functions as a dimer, in which the CTD^HSP90^ forms a relatively stable dimerization interface (30). The NTD^HSP90^ alternates between the open and closed conformations in response to the ATPase cycle (30). To stabilize the HSP90 dimer during purification, molybdate was added in the buffer, which is known to trap HSP90 in the closed state that corresponds to its client-bound conformation (31–33). The PINK1–HSP90β–CDC37 complex was first purified by affinity chromatography (Supplementary Fig. 1a), followed by gel filtration chromatography (Supplementary Fig. 1b). The subunit composition of the complex was confirmed by SDS–PAGE (Supplementary Fig. 1b). Fractions corresponding to the peak top of the gel-filtration profile containing PINK1, HSP90β, and CDC37 were collected and rapidly vitrified on EM grids for single-particle cryo-EM analysis.

### Cryo-EM single particle analysis of the human PINK1–HSP90–CDC37 complex in native conditions

Among 1,606,203 particles picked from 4,217 movies without templates with particle diameters ranging from 100 to 200 Å, 1,373,615 particle images were extracted with a box size of 400 pixels (0.724 Å/pixel) and downsampled to a size of 96 pixels (3.02 Å/pixel) by Fourier cropping (Supplementary Fig. 2). The downsampled images were subjected to reference-free 2D classification, which generated the dimeric projection images. 62 classes with 758,357 particles were used as templates for picking particles with a particle diameter of 200 Å. 1,586,437 particle images were extracted and subjected to the second round of 2D classification, resulting in a set of images similar to that generated in the first round of 2D classification. We selected 38 classes with clear structural features and used 607,867 particles for the *ab initio* 3D reconstruction. Four classes of *ab initio* 3D models were generated and then refined by heterogeneous refinement. The resultant 3D maps from two classes represented the HSP90 dimer, one in a closed state and the other in a widely open state. For the closed state, 199,855 particles were selected from 208,564 particles after excluding junk particles through 2D classification and performed homogeneous refinement and non-uniform refinement.

Non-uniform refinement yielded a density map at a nominal resolution of 3.05 Å. For the widely open state, 211,262 particles were selected from 216,240 particles after 2D classification and performed homogeneous refinement and non-uniform refinement. Non-uniform refinement yielded a density map at a nominal resolution of 3.04 Å.

The cryo-EM analysis revealed two distinct states of the PINK1–HSP90β–CDC37 complex. One state indicates that the HSP90 dimer adopts a closed conformation. In this state, weak density corresponding to the client is visible. The absence of the double-helix of CDC37 suggests that CDC37 has dissociated. In the other state, the HSP90 dimer displays an unusually open conformation at the CTD^HSP90^. The density corresponding to both PINK1 and CDC37 are not observed. While conformational opening and closing of the NTD^HSP90^ are well established for HSP90 (34), such a pronounced separation of the CTD^HSP90^ has not previously been reported. During grid preparation, contact with strong forces at the air–water interface is known to dissociate the protein complex, leading to substantial grid-induced compositional heterogeneity (35). Considering the possibility that the complex was disassembled during vitrification, we next attempted chemical crosslinking to stabilize the complex.

### Cryo-EM single particle analysis of the human PINK1–HSP90–CDC37 complex with chemical crosslinking

To stabilize PINK1 in complex with HSP90 and CDC37, the eluate from the affinity chromatography was crosslinked with glutaraldehyde at a final concentration of 0.025% (Supplementary Fig. 3a), and further purified by size-exclusion chromatography (Supplementary Fig. 3b). SDS–PAGE analysis confirmed the integrity of the crosslinked complex (Supplementary Fig. 3b), which eluted at a position nearly identical to that of the native complex. Fractions corresponding to the peak were collected and rapidly vitrified on EM grids for cryo-EM analysis.

Among 1,610,071 particles picked from 4,281 movies without templates with particle diameters ranging from 100 to 200 Å, 1,379,120 particle images were extracted with a box size of 400 pixels (0.724 Å/pixel) and downsampled to a size of 96 pixels (3.02 Å/pixel) by Fourier cropping (Supplementary Fig. 4a). The downsampled images were subjected to reference-free 2D classification, which generated the dimeric projection images. 95 classes with 1,325,908 particles were used as templates for picking particles with a particle diameter of 200 Å. 1,605,673 particle images were extracted and subjected to the second round of 2D classification, resulting in a set of images similar to that generated in the first round of 2D classification. We selected 120 classes with clear structural features and used 1,368,585 particles for the *ab initio* 3D reconstruction. Four classes of *ab initio* 3D models were generated in the first round and refined by heterogeneous refinement. One class comprising 474,125 particles was further subjected to a second round of *ab initio* reconstruction into three classes followed by heterogeneous refinement. After excluding junk particles through 2D classification, 227,582 particles were selected from the 233,619 particles. Homogeneous and non-uniform refinements were performed to generate density maps. To further optimize the map quality, global and local CTF refinements were applied after non-uniform refinement. The subsequent homogeneous refinement, non-uniform refinement, and two rounds of local refinement yielded a 3.20 Å map. Although the second round of local refinement used the map from the first round as a reference to further optimize particle alignment, the resolution showed little improvement. To improve the accuracy of particle identification, we performed particle picking using the neural network–based method Topaz and obtained 1,171,684 particle images. After 2D classification to remove junk particles, 1,163,406 particles were used for *ab initio* 3D reconstruction and heterogeneous refinement. A single class of 454,727 particles was subjected to homogeneous, non-uniform, and local refinements, resulting in a 3.08 Å map. The atomic models of the HSP90–CDC37 complex (PDB 7Z38) and the predicted PINK1 C-lobe (AlphaFold model, AF-Q9BXM7-F1-model_v4) fit well into the density (local resolution range of 2.5 to 8.5 Å) (Supplementary Figs. 4a, b and Table 1). When we built an atomic model of the complex, no experimentally determined structure of the C-lobe of human PINK1 was available. Therefore, we used the AlphaFold-predicted model.

**Table 1.** Data collection and refinement statistics.

| **PINK1–HSP90–CDC37 complex** |  |
| --- | --- |
| **Data collection and processing** |  |
| Magnification | 190,000 |
| Voltage (kV) | 200 |
| Electron exposure (e<sup>-</sup>/Å<sup>2</sup>) | 50.0 |
| Defocus range (μm) | -0.8 – -1.6 |
| Pixel size (Å) | 0.724 |
| Symmetry imposed | C1 |
| Initial particle images (no.) | 1,610,071 |
| Final particle images (no.) | 454,727 |
| Map resolution (Å) | 3.08 |
| FSC threshold | 0.143 |
| **Refinement** |  |
| Initial model used | 7Z38, AF-Q9BXM7-F1-model_v4 |
| Model resolution (Å) |  |
| FSC 0.143, unmasked/masked | 3.06/3.19 |
| Model composition |  |
| Non-hydrogen atoms | 14,094 |
| Protein residues | 1,732 |
| Ligand | 4 |
| B factors (Å<sup>2</sup>), min/max/mean |  |
| Protein | 80.30/370.77/177.18 |
| Ligand | 88.67/97.64/90.87 |
| RMS deviation |  |
| Bond length (Å) | 0.006 |
| Bond angle (°) | 0.826 |
| Molprobity score | 1.97 |
| Clash score | 16.40 |
| Rotamer outliers (%) | 0.00 |
| Ramachandran plot |  |
| Favored (%) | 96.20 |
| Allowed (%) | 3.80 |
| Disallowed (%) | 0.00 |

### Overall structure of the human PINK1–HSP90–CDC37 complex

Cryo-EM density of the closed HSP90 dimer was observed along with protrusions corresponding to the two α-helices of the NTD^CDC37^ (Fig. 1c). The CDC37 linker was traced extending across the outer surface of HSP90, with weak density attributable to the middle domain. Residual density observed near the complex corresponded well to the known position of the C-lobe of a kinase bound to the HSP90–CDC37 complex. The AlphaFold2-predicted model of PINK1 comprising the C-lobe and CTE (residues 318–581) was fitted into the observed density, although no corresponding density was visible for a part of the activation loop. Additional density was also observed inside the channel formed by the HSP90 dimer, which appeared to be connected to the PINK1 C-lobe.

### Interaction between the C-lobe of PINK1 and the NTD^CDC37^ mediated by the HPNI motif

The density at the interface between the C-lobe of PINK1 and the NTD^CDC37^ is well resolved (Figs. 2a, b). The C-lobe of PINK1 interacts with the NTD^CDC37^ through hydrophobic residues (Figs. 2c-e). The region from His20 to Trp31 of CDC37 are buried within the PINK1 C-lobe (Figs. 2c-e). Trp31 of CDC37 is positioned in close proximity to Leu353, Asp384 of the DFG motif, and Asp360 of the HRD motif of PINK1 (Fig. 2c). Ile23 of CDC37 forms hydrophobic interactions with Ile382 in β7 and Leu353 of the αE helix of PINK1 (Fig. 2d). His352 of PINK1 forms a π-π stacking interaction with His20 of CDC37 (Fig. 2e). The PINK1 residues forming the CDC37-interacting interface are highly conserved across vertebrate orthologs (Fig. 2f), highlighting the functional importance of this interface for the recognition of PINK1 by CDC37.

**Figure 2.**
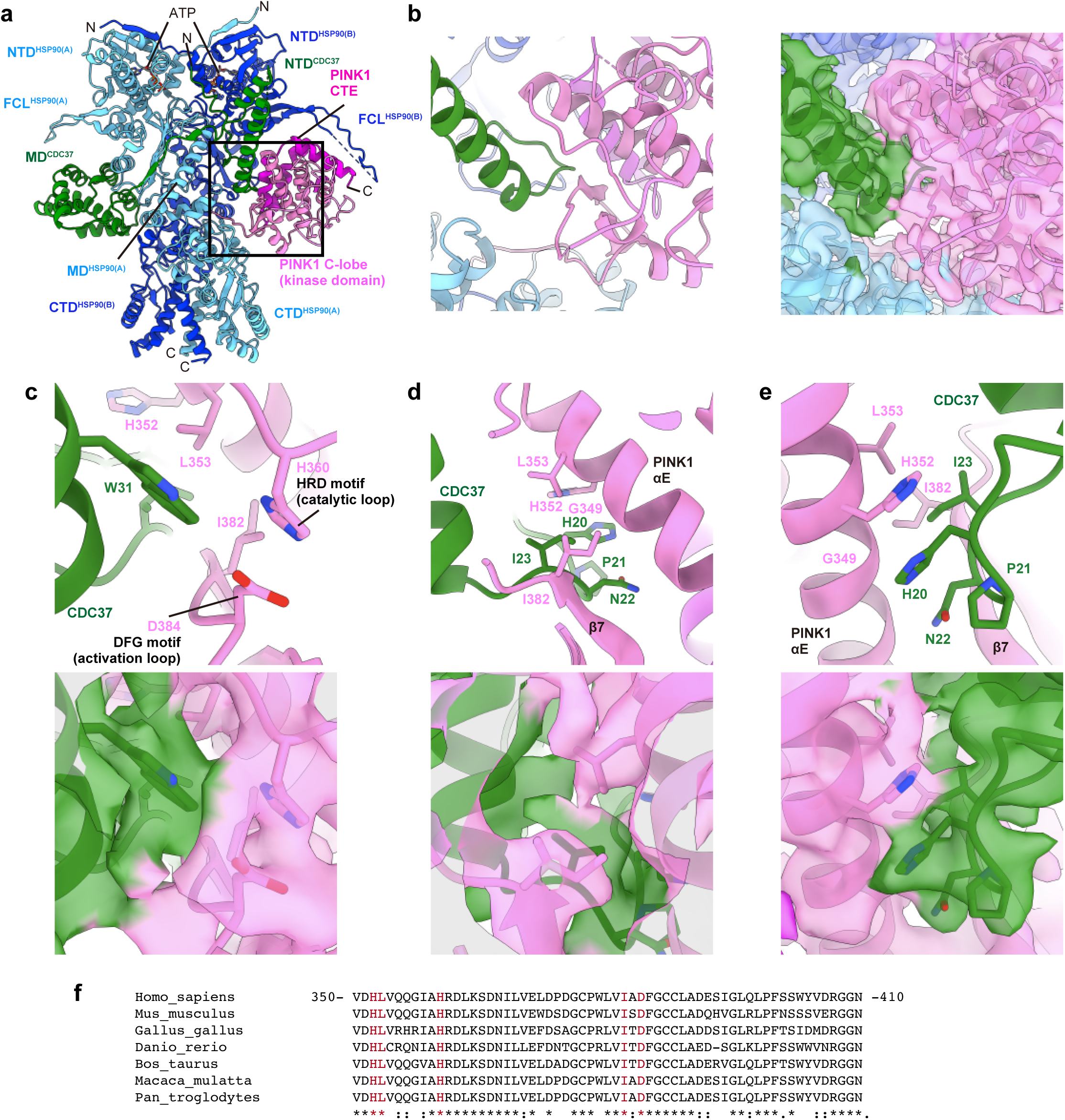
Interaction of the PINK1 C-lobe with CDC37 mediated by the loop of the N-terminal domain. **(a)** Overall view of the cryo-EM structure of the PINK1–HSP90–CDC37 complex. The box indicates an area including the locations of the views shown in (b-e) **(b)** Close-up view of the density map at the interface between PINK1 (pink) and CDC37 (green). **(c)** Close-up view of the PINK1–CDC37 interaction around Trp31 of CDC37. The structure (top) and corresponding map (bottom) are shown. **(d)** Close-up view of the PINK1–CDC37 interaction around Ile23 of CDC37. The structure (top) and corresponding map (bottom) are shown. **(e)** Close-up view of the PINK1–CDC37 interaction around His31 of CDC37. The structure (top) and corresponding map (bottom) are shown. **(f)** Amino-acid sequence alignment of the PINK1 residues located at the CDC37-interacting interface among representative vertebrates. *, identical residues; :, residues with strongly similar properties; ., residues with weakly similar properties.

The loop (residues 20–23) of CDC37 overlaps with the loop (residues 271–274) between insertion2 (ins2) and β4 of the N-lobe of PINK1 in the predicted model (Figs. 3a, b). The primary structures of both loops share the “HPNI” motif (Figs. 3b, c), which is consistent with the established role of this sequence as a critical determinant for the kinase–CDC37 interaction (33, 36). The loop is located on the backside of the ATP-binding pocket and stabilizes the relative positioning of the N- and C-lobes (Fig. 3a). The HPNI motif of PINK1 is evolutionarily conserved (Fig. 3d). The H271Q mutation within this motif in PINK1 has been reported as a Parkinson’s disease-associated mutation (Fig. 3b). These observations suggest that the NTD^CDC37^ associates with PINK1 by mimicking the interactions formed with the folded N-lobe.

**Figure 3.**
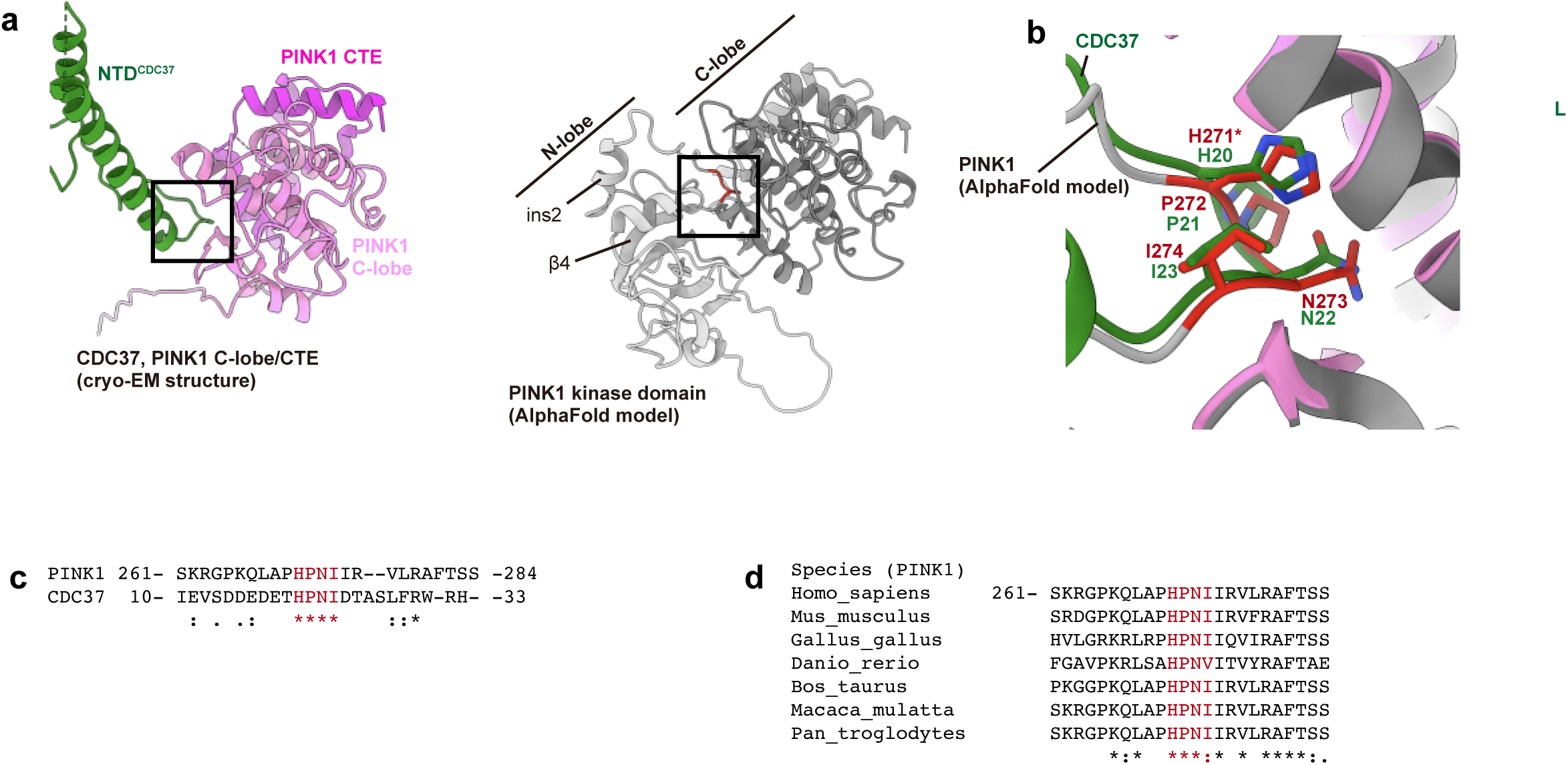
Comparison of the HPNI motif between CDC37 and PINK1. **(a)** Comparison between the NTD^CDC37^–C-lobe^PINK1^ interaction (cryo-EM) and N-lobe^PINK1^–C-lobe^PINK1^ interaction (AlphaFold2). The boxes indicate the loop containing the HPNI motif. **(b)** Close-up view of the HPNI motifs of CDC37 (green; cryo-EM) and PINK1 (red; AlphaFold2). **(c)** Amino-acid sequence alignment of the HPNI motif-containing loop of human PINK1 and CDC37. *, identical residues; :, residues with strongly similar properties; ., residues with weakly similar properties. **(d)** Amino-acid sequence alignment of the HPNI motif-containing loop of PINK1 among representative vertebrates. *, identical residues; :, residues with strongly similar properties; ., residues with weakly similar properties.

### The β5 strand of the PINK1 N-lobe passing through the HSP90 dimer channel

The five β strands of the PINK1 N-lobe are a conserved structural element that contributes to the formation of the ATP-binding pocket, together with the C-lobe. The β5 strand of PINK1 is located immediately upstream of the C-lobe (Fig. 4a). Within the central channel of the HSP90 dimer, we observed residual density corresponding to the β5 strand of PINK1 and extending N-terminally up to Leu308 (Figs. 4b-e). The hydrophobic side chains of Phe315, Leu316, and Val317, on β5 of PINK1 are buried in a groove formed by Leu439, Ile517, Tyr520, and Leu611 in one of the two HSP90 protomers (HSP90^A^) (Figs. 4d, e). On the other hand, no density could be clearly assigned to the other four β strands of the PINK1 N-lobe (Figs. 1c, 4c). These findings suggest that the antiparallel β sheet of the PINK1 N-lobe could not assemble in the PINK1–HSP90–CDC37 complex, resulting in unfolding of the N-lobe. The HSP90 dimer captures PINK1 in an intermediate folding state, where the C-lobe and CTE is folded but the N-lobe is unfolded.

**Figure 4.**
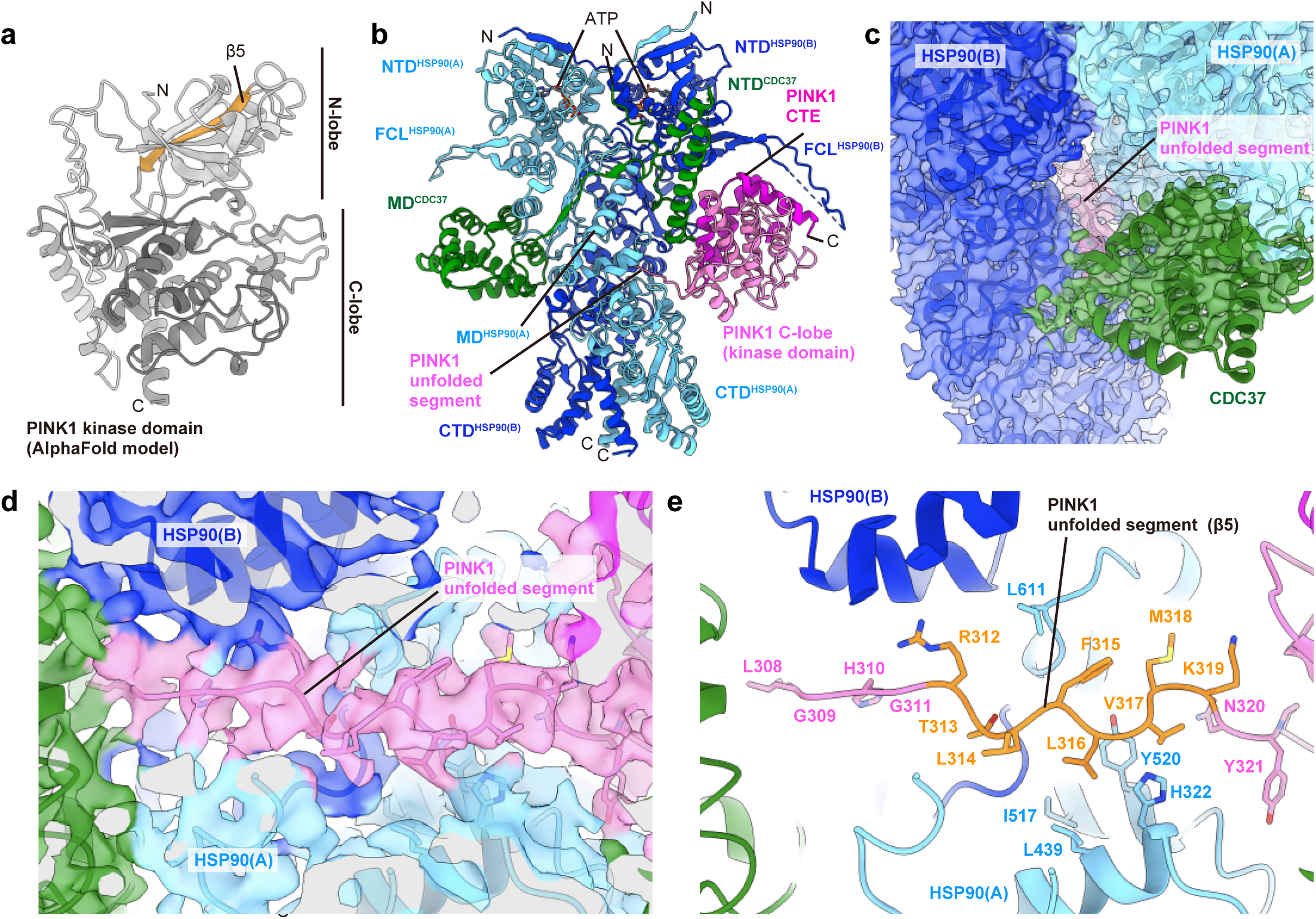
Insertion of the β5 strand of the PINK1 N-lobe into the interfacial channel of the HSP90 dimer. **(a)** The AlphaFold2-predicted model (AF-Q9BXM7-F1-model_v4) of human PINK1. The N-lobe contains the five-stranded β sheet (β1–β5). (**b)** Overall view of the cryo-EM structure of the PINK1–HSP90–CDC37 complex. **(c)** Close-up view of the unfolded segment of the PINK1 N-lobe, which is inserted into the channel at the interface between two protomers in the HSP90 dimer. **(d, e)** Close-up view of the density map and structures around the unfolded segment of the PINK1 N-lobe.

### Accommodation of the PINK1 C-lobe and CTE in the FCL–MD space of the HSP90 dimer

The CTE of PINK1 is composed of a short helix (αJ) followed by three α-helices (αK–αM) connected by the loops, which is characteristic of a unique structural element not found in conventional kinases. In the present complex structure, the CTE of PINK1 is folded and positioned back onto the kinase domain (Figs. 1c, 5a). The C-lobe and CTE of PINK1 are accommodated in the cleft formed by the MD^HSP90^ and FCL^HSP90^ (Figs. 1c, 5a). The position of the αK helix in the AlphaFold2 model of PINK1 would clash with HSP90 (Fig. 5b). The αK helix is suggested to rotate to avoid the clash with HSP90. Leu539 on the αK helix of PINK1 hydrophobically interacts with Trp312 of HSP90 (Fig. 5c). The αJ–αK linker and αM helix of PINK1 are positioned close to the FCL^HSP90^ (Fig. 5d). Due to discontinuous density, the atomic model of the FCL^HSP90^ could not be built (Fig. 5d). The interaction between PINK1 and the FCL^HSP90^ appears to be relatively weak. To explore the conformational heterogeneity of the FCL^HSP90^, we performed 3D Variability Analysis (3D VA) in CryoSPARC. This analysis visualized relative motions of the FCL^HSP90^ to the CTE of PINK1 in accordance with lower local resolution in the periphery of the complex (Supplementary Fig. 5). These structural features indicate that the FCL^HSP90^ is loosely or not bound to the CTE of PINK1. Because the sample preparation involved chemical crosslinking to stabilize this dynamic complex, we cannot fully exclude the possibility that crosslinking affected the relative configuration of these flexible elements. Nonetheless, the continuous motions revealed by 3D VA support an intrinsically dynamic interface.

**Figure 5.**
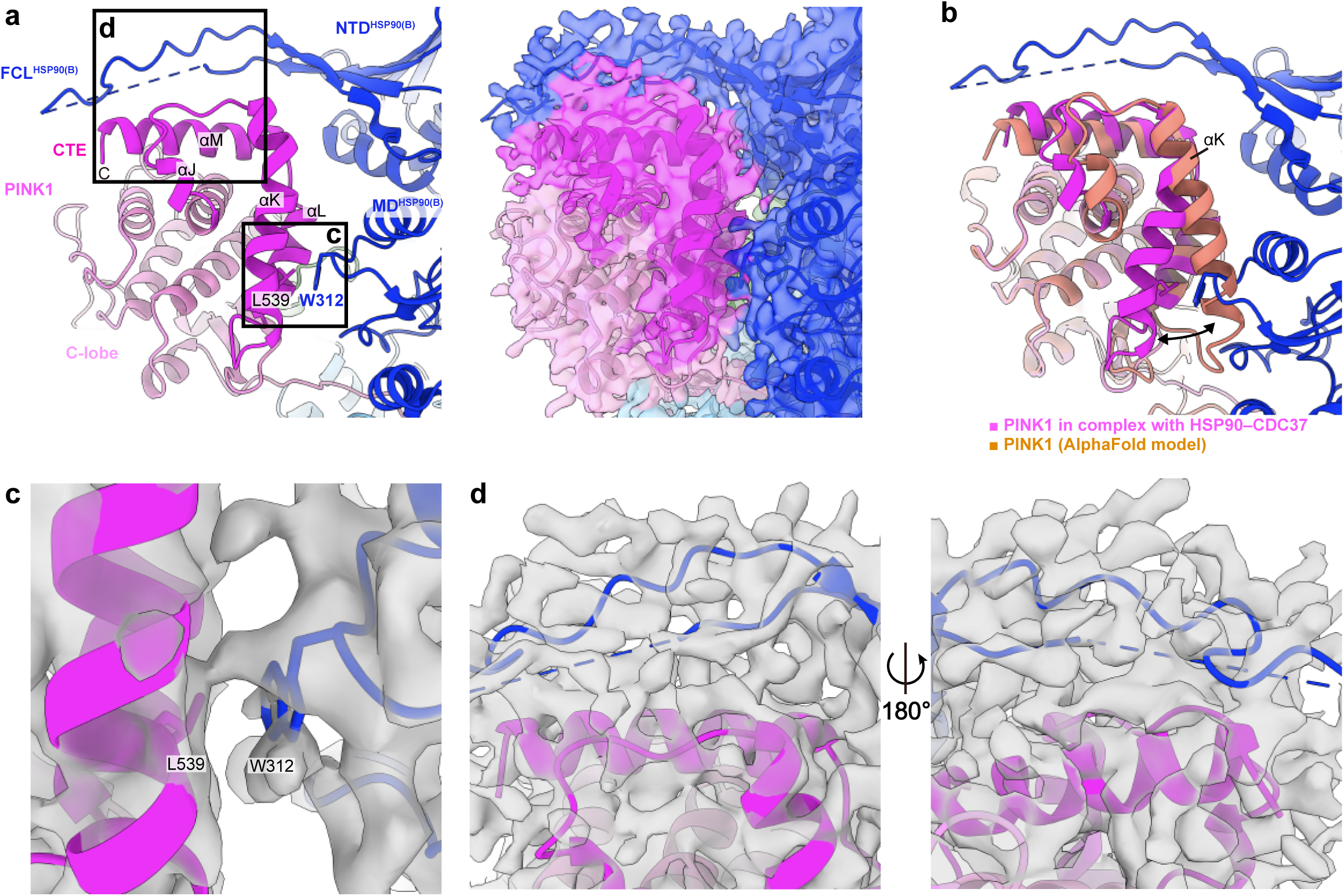
Interface between the PINK1 CTE and HSP90. **(a)** The structure and density map of the interface between PINK1 and HSP90. The two boxes indicate the locations of the views shown in (c) and (d) **(b)** Superposition of the cryo-EM and AlphaFold2-predicted PINK1 structures. **(c)** Close-up view of the structure and density map around Trp312 of HSP90 and Leu539 of PINK1. **(d)** Close-up view of the interface between the PINK1 CTE and HSP90 FCL.

### Overlapping surfaces around the CTE of PINK1 with HSP90 and the TOM complex

PINK1 forms a complex with the translocase of the outer membrane (TOM) and VDAC on the outer membrane of depolarized mitochondria (20). In the PINK1–TOM–VDAC complex, the CTE of PINK1 interacts with the N-helix of PINK1 and the TOM20 and TOM5 components of the TOM complex (Figs. 6a, b). The CTE of PINK1 faces the MD^HSP90^ and FCL^HSP90^ in the present PINK1–chaperone complex structure. To clarify the difference in the interaction of the PINK1 CTE with HSP90 and the TOM complex, we aligned and superposed the structures of the HSP90–CDC37 and TOM–VDAC complexes, using the PINK1 CTE as the reference (Figs. 6c, d). The N-helix of PINK1 makes intramolecular contacts with the αK and αL helices. In the TOM–VDAC complex, TOM20 interacts with the N-helix and αK helix of PINK1, whereas TOM5 interacts with the αM helix of PINK1. The positions of the N-helix of PINK1 and TOM20 partially overlap with that of the MD^HSP90^ (Fig. 6c). The position of TOM5 overlaps with that of the FCL^HSP90^ (Fig. 6d). These findings suggest that the interactions of PINK1 with the HSP90–CDC37 and TOM complexes are competitive and mutually exclusive.

**Figure 6.**
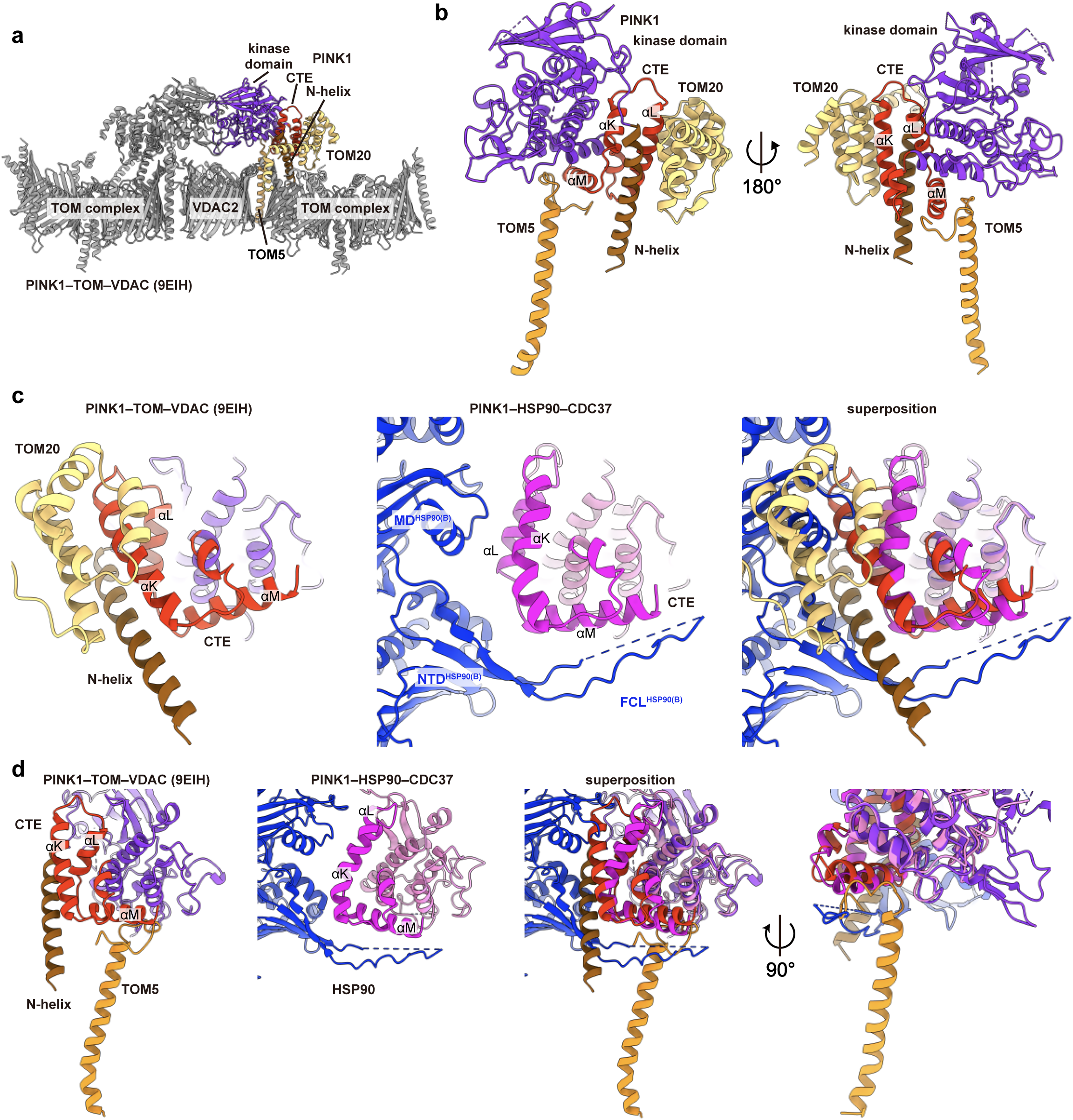
Comparison of the spatial arrangement of HSP90–CDC37 and TOM–VDAC relative to PINK1. **(a)** Cryo-EM structure of the PINK1–TOM–VDAC complex (PDB 9EIH). **(b)** Close-up views of PINK1–TOM5–TOM20 in the PINK1–TOM–VDAC complex. The other components of the complex are omitted for clarity. **(c)** Comparison between the PINK1–TOM20 and PINK1–HSP90 interactions. In the PINK1–TOM20 interface (left panel), TOM20 interacts with both the PINK1 CTE and N-helix, which also interact with each other. In the PINK1–HSP90 interface (middle panel), the FCL^HSP90^ covers the PINK1 CTE. The superposition (right panel) shows that the PINK1 N-helix and TOM20 overlap with the NTD^HSP90^ and MD^HSP90^. **(d)** Comparison between the PINK1–TOM5 and PINK1–HSP90 interactions. In the PINK1–TOM5 interface (left panel), the TOM5 interacts with the PINK1 CTE. The superposition (right panel) shows that TOM5 overlaps with the FCL^HSP90^.

## Discussion

PINK1 has been identified as a high-affinity client of the HSP90–CDC37 chaperone complex by a kinome-wide quantitative screen and mass spectrometry (11, 23, 27, 28). HSP90 inhibitors such as 17-AAG and geldanamycin decrease the half-life and protein levels of PINK1 under normal conditions (28, 29). On the other hand, when mitochondrial depolarization and HSP90 inhibition occur simultaneously, different cellular dynamics of PINK1 have been observed: the amount of PINK1 protein and ubiquitin phosphorylation are decreased in some studies (37, 38), whereas the half-life of PINK1 is prolonged and its ability to recruit Parkin remains, regardless of HSP90 inhibition, in other studies (29, 39). Thus, the role of HSP90 in regulating PINK1 under mitochondrial depolarization conditions remains controversial. Nonetheless, there is a consensus that PINK1 stability is strongly influenced by the HSP90 chaperone system under normal conditions. In this study, we determined the cryo-EM structure of the human PINK1–HSP90–CDC37 complex. In the present structure, the C-lobe and the CTE of PINK1 are already folded, whereas the N-lobe of PINK1 likely remains unfolded. This structure shows that the HSP90–CDC37 complex holds a folding intermediate of PINK1, which stabilizes key structural elements prior to full kinase domain maturation. The intermediate state suggests a stepwise folding mechanism of the kinase domain of PINK1.

CDC37 functions as a kinase-specific co-chaperone of HSP90, which associates with immature kinases and facilitates their delivery to HSP90 (25). The HPNI amino acids sequence of CDC37 is the critical conserved motif for kinase recognition (36). Disruption of this motif by mutation reduces binding affinity for client kinases, thereby impairing their stability and maturation in the cell (33, 40). The present cryo-EM structure of the PINK1–HSP90–CDC37 complex shows that the HPNI motif of CDC37 also interacts with the C-lobe of PINK1 in a manner similar to the HPNI motif of PINK1 in the AlphaFold2-predicted PINK1 structure.

The L347P and H271Q mutations of PINK1 have been identified in patients with Parkinson’s disease (41, 42). The L347P mutation decreases the stability of PINK1 due to impaired interaction with the HSP90–CDC37 complex (26, 28). In the present structure, Leu347 of PINK1 is located on the αE helix of the C-lobe, which interacts with the HPNI motif of CDC37. The L347P mutation may distort αE and disrupt its interaction with CDC37. On the other hand, the H271Q mutation is located within the HPNI motif of the N-lobe of PINK1. The H271Q mutation potentially interferes with the π–π stacking between His271 and His352. We previously reported that the H271Q mutant of PINK1 is defective in complex formation and autophosphorylation upon mitochondrial depolarization, thereby impairing the recruitment and activation of Parkin on damaged mitochondria (4, 10, 13). The H271Q mutation is suggested to affect the integrity of the kinase domain through destabilization of the interaction between the N-lobe and C-lobe, which likely hinders proper folding. Our findings provide the molecular basis for familial Parkinson’s disease linked to PINK1 mutations.

A recent cryo-EM study has revealed the structure of a PINK1–TOM–VDAC array that promotes PINK1 stagnation to mitochondria and activation through trans-autophosphorylation (20). The CTE of PINK1 interacts with the N-helix and with TOM5 and TOM20, components of the TOM complex. The present PINK1–HSP90–CDC37 structure shows that the HSP90–CDC37 complex shields the CTE of PINK1. As the position of the HSP90–CDC37 complex relative to the CTE of PINK1 overlaps with those of TOM5 and TOM20 in the PINK1–TOM–VDAC array, the HSP90–CDC37 complex is likely to dissociates from PINK1 when PINK1 is recruited to damaged mitochondria. Release from the HSP90–CDC37 complex would enable folding of the N-lobe of PINK1 and intramolecular interactions between the N-helix and CTE of PINK1, thereby facilitating maturation of the kinase domain. In addition, the CTE of PINK1 can interact with components of the TOM complex, which in turn permits the activation of PINK1 through trans-autophosphorylation. These findings suggest that the interactions of PINK1 with the HSP90–CDC37 complex and with the TOM complex are mutually exclusive and that the HSP90–CDC37 complex could not promote the activation of PINK1 cooperatively with the TOM complex. Diverse effects of the HSP90 inhibition on the cellular dynamics of PINK1 may be due to a role of the HSP90–CDC37 complex, which is stabilizing PINK1 in an intermediate folding state and does not directly influence the PINK1 activation upon mitochondrial depolarization.

Here, we propose a model in which PINK1 is held in an intermediate conformation by the HSP90–CDC37 complex and that the kinase domain of PINK1 completes folding through interactions with the TOM complex and dissociation from the HSP90–CDC37 complex on the outer membrane of damaged mitochondria (Fig. 7). The HSP90–CDC37 system involves other co-chaperones and post-translational modifications. The phosphorylation of CDC37 at Ser13 by casein kinase 2 (CK2) has been shown to be a prerequisite for the interaction between CDC37 and client kinases (43, 44). In contrast, protein phosphatase 5 (PP5) promotes dissociation of client kinases from the HSP90–CDC37 complex by dephosphorylating CDC37 at Ser13 (44, 45). In addition, activator of HSP90 ATPase activity 1 (Aha1) is a co-chaperone known to stimulate the ATP hydrolysis of HSP90 (46, 47). Further studies are required to understand that co-chaperones and post-translational modifications of the HSP90–CDC37 complex fine-tune the PINK1 stability and activity. Our results provide a structural basis for PINK1 stabilization and reveal a mechanism by which the HSP90–CDC37 complex facilitates PINK1 folding and protects its unengaged interface.

**Figure 7.**
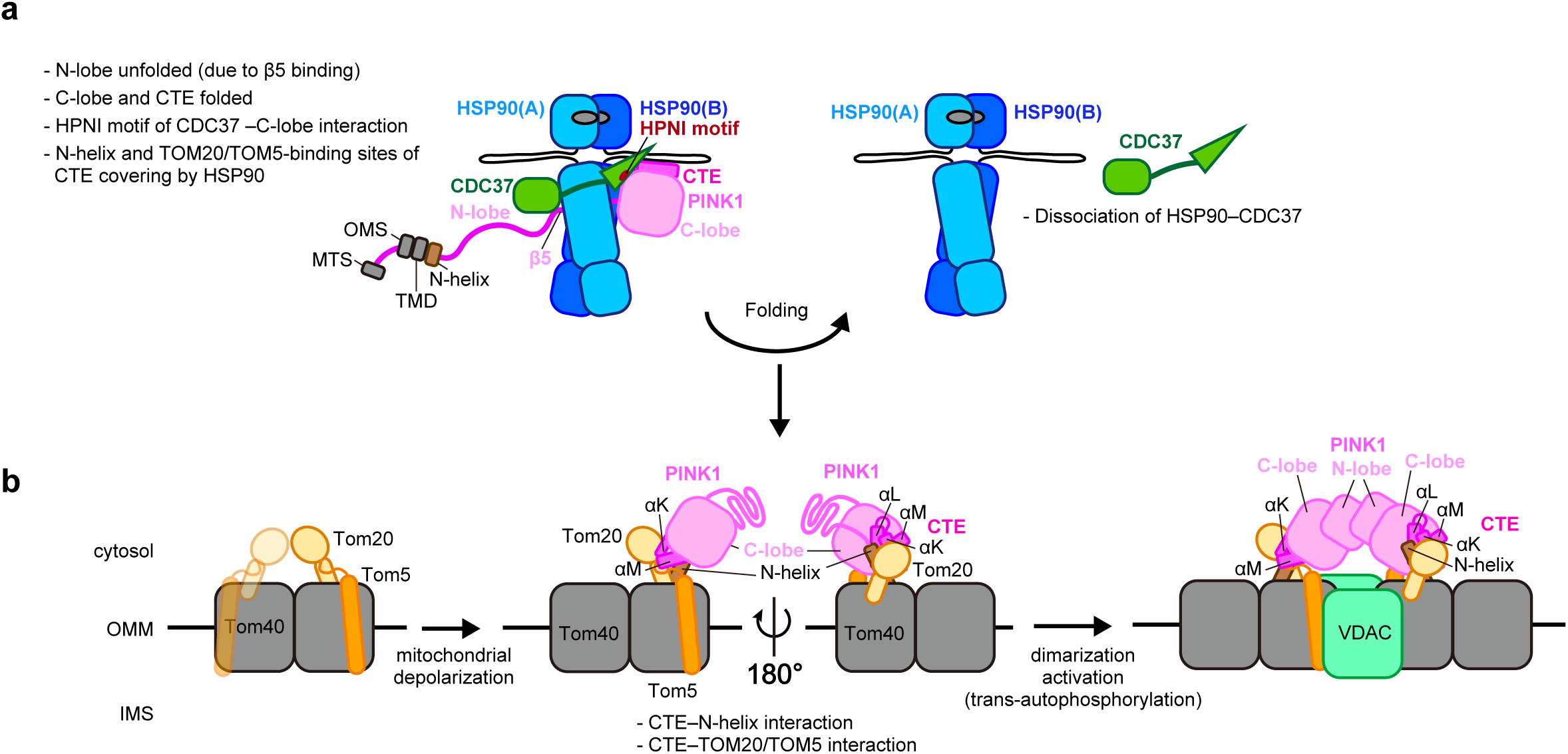
Model for PINK1 stabilization by the HSP90–CDC37 complex and transition to its active state on depolarized mitochondria. **(a)** In the cytosol, the HSP90–CDC37 complex captures the folding intermediate of PINK1, where the N-lobe remains unfolded due to binding of the isolated β5 strand to HSP90, whereas the C-lobe and CTE is folded. The CTE is covered with FCL^HSP90^. The HPNI motif of CDC37 interacts with the C-lobe of PINK1, mimicking the HPNI motif of the N-lobe. **(b)** TOM20 and TOM5 compete with HSP90–CDC37 for binding to the PINK1 CTE, displacing the chaperone complex from PINK1. **(c)** Upon mitochondrial depolarization, the dissociation of the HSP90–CDC37 complex allows the PINK1 CTE to interact with the N-helix and further with TOM20 and TOM5, along with the N-lobe folding. Finally, PINK1 dimerizes and becomes activated in a trans-autophosphorylation fashion.

## Materials and methods

### Plasmids

Human PINK1 (residues 104–581, accession No. NM_032409.3), in which Phe104 was replaced by Met, was cloned with a C-terminal His₈ tag into the BamHI and EcoRI sites downstream of the PH promoter in the pFastBacDual vector. Human HSP90β (accession No. NM_001271969.2) and human CDC37 (accession No. NM_007065.4) were cloned into the BamHI and EcoRI sites under the PH promoter and into the KpnI and XhoI sites under the P10 promoter, respectively.

### Sample preparation for cryo-EM

PINK1(F104M)-His₈, HSP90β, and CDC37 were co-expressed in Sf9 cells using the Bac-to-Bac system (Invitrogen) with X-tremeGENE HP (Roche) as the transfection reagent for P1 virus generation. The P2 virus was amplified to produce 20 mL of P3 virus, which was subsequently used to infect 200 mL of Sf9 cells. The infected cells were incubated at 27 °C for 3 days. The infected Sf9 cells were harvested by centrifugation at 2,000 × g for 5 min at 4 °C and washed twice with cold phosphate buffered saline (PBS). The cell pellets were resuspended in a lysis buffer (50 mM Tris-HCl, pH 8.0, 150 mM NaCl, 10 mM MgCl₂, 10 mM KCl, 20 mM Na₂MoO₄, 20 mM imidazole, and 0.5% Triton X-100) and lysed by sonication on ice for 5 min. The lysate was clarified by centrifugation at 2,000 × g for 10 min at 4 °C. The cleared lysate was loaded onto a Ni-NTA agarose (Qiagen) column pre-equilibrated with the lysis buffer. The column was washed with the lysis buffer (first wash), without Triton X-100 (second wash). The bound protein was eluted with 50 mM Tris-HCl buffer (pH 8.0) containing 150 mM NaCl and 10 mM MgCl_2_, 10 mM KCl, 20 mM Na2MoO_4_, and 300 mM imidazole. For purification of the native complex, the elution sample was further purified by a size-exclusion chromatography on a ENrich™ SEC 650 column (10 × 300 mm; Bio-Rad) equilibrated with 50 mM Tris-HCl buffer (pH 8.0) containing 150 mM NaCl and 10 mM MgCl_2_, 10 mM KCl, and 20 mM Na_2_MoO_4_. A fraction enriched in PINK1 (F104M)-His₈, HSP90β, and CDC37 was concentrated to 0.67 g/L using an Amicon Ultra-4 centrifugal filter unit (100 kDa MWCO) for cryo-EM analysis. For preparation of the crosslinked complex, the elution sample was concentrated tenfold and diluted three times using an Amicon Ultra-4 centrifugal filter unit (100 kDa MWCO), thereby exchanging the buffer to the crosslinking buffer (50 mM HEPES-Na, pH 8.0, 150 mM NaCl, 10 mM MgCl₂, 10 mM KCl, and 20 mM Na₂MoO₄). Crosslinking was performed by adding glutaraldehyde at a final concentration of 0.025% and incubating at room temperature for 30 min. The reaction was quenched with Tris-HCl buffer (pH 8.0) at a final concentration of 0.1 M. The crosslinked sample was further purified by size-exclusion chromatography on an ENrich™ SEC 650 column (10 × 300 mm) equilibrated with 50 mM Tris-HCl (pH 8.0), 150 mM NaCl, 10 mM MgCl₂, 10 mM KCl, and 20 mM Na₂MoO₄. The fraction enriched in the crosslinked complex was concentrated to 0.30 g/L using an Amicon Ultra-0.5 centrifugal filter unit (3 kDa MWCO).

### Cryo-EM grid preparation and data collection

Cu grids (R1.2/1.3, 300 mesh, Quantifoil) with holey carbon films were hydrophilized using a JEC-3000FC Auto Fine Coater (JEOL) at 7 Pa, 10 mA, and 10 sec. A 3 μL aliquot of 0.67 g/L intact protein solution or 0.30 g/L crosslinked protein solution was added to the grids using a Vitrobot Mark IV (Thermo Fisher Scientific) at a temperature of 8 °C and 100% humidity. The grids were then immersed in liquid ethane and rapidly frozen under the following conditions: waiting time of 0 sec, blotting time of 5 sec, and blotting force of 15 for the intact sample, and waiting time of 0 sec, blotting time of 3 sec, and blotting force of 15 for the crosslinked sample. Data collection was carried out on a Glacios cryo-transmission electron microscope (Thermo Fisher Scientific) operating at 200 kV, equipped with a Falcon 4 electron detector at Kyoto University. A total of 4,217 or 4,281 movies were automatically collected using EPU software. Movies were collected at a target defocus range of −0.8 to −1.6 μm and a nominal magnification of ×190,000, corresponding to a calibrated pixel size of 0.724 Å/pixel. Each movie was recorded with an exposure time of both 4.31 s, subdivided into both 66 frames with a total electron dose of 50 e^−1^ Å^−2^. All processing was performed in CryoSPARC v.4.3.1, v4.4.1, and v.4.7.0 (48).

### Cryo-EM data processing for the native PINK1–HSP90–CDC37 complex

The collected micrographs were processed using patch motion correction and patch CTF estimation. 1,606,203 particles were initially picked using blob picker (minimum particle diameter: 100 Å, maximum particle diameter: 200 Å). After 2D classification, 758,357 particles were selected based on the average 2D images of the protein and used as templates for template picker (particle diameter: 200 Å). 1,816,520 particles were picked. 607,867 particles selected from the average 2D images of the protein and subjected to *ab initio* reconstruction. Heterogeneous refinement was performed using four 3D references, generated two class with a well-defined structure. 199,855 or 211,262 particles were selected after 2D classification further refined by homogeneous and non-uniform refinement. Local refinement following non-uniform refinement resulted in a 3.05 Å or 3.04 Å resolution 3D map of the PINK1–HSP90–CDC37 complex. Overall resolution estimates correspond to a Fourier shell correlation of 0.143 using an optimized mask that is automatically determined after refinement.

### Cryo-EM data processing for the crosslinked PINK1–HSP90–CDC37 complex

The collected micrographs were processed using patch motion correction and patch CTF estimation. 1,610,071 particles initially were picked using blob picker (minimum particle diameter: 100 Å, maximum particle diameter: 200 Å). 1,325,908 particles were selected from the average 2D images of the protein and used as templates for template picker (particle diameter: 200 Å), resulting in the picked of 1,857,354 particles. 1,368,585 particles were selected from the average 2D images of the protein and subjected to an *ab initio* reconstruction and heterogeneous refinement into four 3D classes. One of the resulting classes was further refined through a second round of *ab initio* reconstruction and heterogeneous refinement into three 3D classes. The selected particles were further cleaned by 2D classification, resulting in 227,582 particles. These particles were processed through homogeneous refinement, non-uniform refinement, global and local CTF refinement, homogeneous refinement, non-uniform refinement, and two rounds of local refinement, yielding a final map at 3.20 Å resolution. Additionally, particle picking was performed using Topaz (49), resulting in 1,171,684 particles. 1,163,406 particles were selected after 2D classification and subjected to *ab initio* reconstruction into four classes, followed by heterogeneous refinement. 454,727 particles were selected and refined by homogeneous and non-uniform refinement. Local refinement following non-uniform refinement resulted in a 3.08 Å resolution 3D map of the PINK1–HSP90–CDC37 complex. Overall resolution estimates correspond to a Fourier shell correlation of 0.143 using an optimized mask that is automatically determined after refinement. Local resolution maps were obtained using local resolution estimation. 3D variability analysis (50) was performed on the final 454,727 particle set, using a mask that encompassed the PINK1, solving for 3 orthogonal principal modes over 20 iterations with a 8 Å resolution filter. The resulting variability was visualized using 3D variability display in cluster mode, with the number of frames/clusters set to six.

### Model building

Model building was performed using Coot (51) and UCSF ChimeraX (52). The initial model of the HSP90–CDC37 complex used a part of the cryo-EM structure of the HSP90–CDC37 complex (PDB 7Z38) (33). The initial model of the C-lobe and CTE of PINK1 used AlphaFold model (AF-Q9BXM7-F1-model_v4). These models fitted to the cryo-EM map using UCSF ChimeraX. The final structure was refined using Phenix (53). All figures were created using UCSF ChimeraX.

## Data availability

The coordinates and maps of the PINK1–HSP90–CDC37 complex have been deposited in the Protein Data Bank/Electron Microscopy Data Bank under the accession codes of 9X4X/EMD-66567. Other data are available from the corresponding authors upon reasonable request.

## Acknowledgements

Cryo-EM data collection was performed using a Glacios cryo-TEM with the support of the Cryo-EM Facility, Institute for Life and Medical Sciences, Kyoto University. Joint Usage/Research Center Program of the Institute for Life and Medical Sciences at Kyoto University (to T.N.). This work was supported by JSPS/MEXT KAKENHI (21K15084, 24H01894 to K.O. and 18H05501 to S.F.)

## Author contributions

K.O.-Conceptualization, Data curation, Formal analysis, Funding acquisition, Investigation, Methodology, Project administration, Resources, Software, Visualization, Writing – original draft.

H.Y.-Investigation, Writing – review and editing.

A.O.-Investigation, Writing – review and editing.

S.G.-Investigation, Resources

Y.N.-Investigation, Resources.

Y.S.-Resources, Supervision

T.N.-Resources, Supervision

S.F.-Supervision, Funding acquisition, Software, Project administration, Writing – review and editing.

## Additional information

We declare no competing financial interests.

## Notes

### Competing Interest Statement

The authors have declared no competing interest.

### Summary of Updates

The main text was edited to improve clarity. Figures were revised to correct labeling and enhance presentation.

## References

1. E. M. Valente et al., PINK1 mutations are associated with sporadic early-onset parkinsonism. Ann Neurol 56, 336–341 (2004).

2. T. Kitada et al., Mutations in the parkin gene cause autosomal recessive juvenile parkinsonism. Nature 392, 605–608 (1998).

3. D. P. Narendra et al., PINK1 is selectively stabilized on impaired mitochondria to activate Parkin. PLoS Biol 8, e1000298 (2010).

4. N. Matsuda et al., PINK1 stabilized by mitochondrial depolarization recruits Parkin to damaged mitochondria and activates latent Parkin for mitophagy. J Cell Biol 189, 211–221 (2010).

5. C. Vives-Bauza et al., PINK1-dependent recruitment of Parkin to mitochondria in mitophagy. Proc Natl Acad Sci U S A 107, 378–383 (2010).

6. S. M. Jin et al., Mitochondrial membrane potential regulates PINK1 import and proteolytic destabilization by PARL. J Cell Biol 191, 933–942 (2010).

7. E. Deas et al., PINK1 cleavage at position A103 by the mitochondrial protease PARL. Hum Mol Genet 20, 867–879 (2011).

8. A. W. Greene et al., Mitochondrial processing peptidase regulates PINK1 processing, import and Parkin recruitment. EMBO Rep 13, 378–385 (2012).

9. C. Zhou et al., The kinase domain of mitochondrial PINK1 faces the cytoplasm. Proc Natl Acad Sci U S A 105, 12022–12027 (2008).

10. K. Okatsu et al., PINK1 autophosphorylation upon membrane potential dissipation is essential for Parkin recruitment to damaged mitochondria. Nat Commun 3, 1016 (2012).

11. K. Yamano, R. J. Youle, PINK1 is degraded through the N-end rule pathway. Autophagy 9, 1758–1769 (2013).

12. M. M. Muqit et al., Altered cleavage and localization of PINK1 to aggresomes in the presence of proteasomal stress. J Neurochem 98, 156–169 (2006).

13. K. Okatsu et al., A dimeric PINK1-containing complex on depolarized mitochondria stimulates Parkin recruitment. J Biol Chem 288, 36372–36384 (2013).

14. C. Kondapalli et al., PINK1 is activated by mitochondrial membrane potential depolarization and stimulates Parkin E3 ligase activity by phosphorylating Serine 65. Open Biol 2, 120080 (2012).

15. M. Iguchi et al., Parkin-catalyzed ubiquitin-ester transfer is triggered by PINK1-dependent phosphorylation. J Biol Chem 288, 22019–22032 (2013).

16. F. Koyano et al., Ubiquitin is phosphorylated by PINK1 to activate parkin. Nature 510, 162–166 (2014).

17. A. F. Schubert et al., Structure of PINK1 in complex with its substrate ubiquitin. Nature 552, 51–56 (2017).

18. C. Gladkova, S. L. Maslen, J. M. Skehel, D. Komander, Mechanism of parkin activation by PINK1. Nature 559, 410–414 (2018).

19. S. A. Sarraf et al., Landscape of the PARKIN-dependent ubiquitylome in response to mitochondrial depolarization. Nature 496, 372–376 (2013).

20. S. Callegari et al., Structure of human PINK1 at a mitochondrial TOM-VDAC array. Science 388, 303–310 (2025).

21. O. G. Raimi et al., Mechanism of human PINK1 activation at the TOM complex in a reconstituted system. Sci Adv 10, eadn7191 (2024).

22. M. A. Eldeeb et al., Tom20 gates PINK1 activity and mediates its tethering of the TOM and TIM23 translocases upon mitochondrial stress. Proc Natl Acad Sci U S A 121, e2313540121 (2024).

23. M. Taipale et al., Quantitative analysis of HSP90-client interactions reveals principles of substrate recognition. Cell 150, 987–1001 (2012).

24. L. Stepanova, X. Leng, S. B. Parker, J. W. Harper, Mammalian p50Cdc37 is a protein kinase-targeting subunit of Hsp90 that binds and stabilizes Cdk4. Genes Dev 10, 1491–1502 (1996).

25. Y. Kimura et al., Cdc37 is a molecular chaperone with specific functions in signal transduction. Genes Dev 11, 1775–1785 (1997).

26. W. Lin, U. J. Kang, Structural determinants of PINK1 topology and dual subcellular distribution. BMC Cell Biol 11, 90 (2010).

27. A. Weihofen, B. Ostaszewski, Y. Minami, D. J. Selkoe, Pink1 Parkinson mutations, the Cdc37/Hsp90 chaperones and Parkin all influence the maturation or subcellular distribution of Pink1. Hum Mol Genet 17, 602–616 (2008).

28. Y. Moriwaki et al., L347P PINK1 mutant that fails to bind to Hsp90/Cdc37 chaperones is rapidly degraded in a proteasome-dependent manner. Neurosci Res 61, 43–48 (2008).

29. W. Lin, U. J. Kang, Characterization of PINK1 processing, stability, and subcellular localization. J Neurochem 106, 464–474 (2008).

30. C. Prodromou et al., The ATPase cycle of Hsp90 drives a molecular ’clamp’ via transient dimerization of the N-terminal domains. EMBO J 19, 4383–4392 (2000).

31. S. D. Hartson, V. Thulasiraman, W. Huang, L. Whitesell, R. L. Matts, Molybdate inhibits hsp90, induces structural changes in its C-terminal domain, and alters its interactions with substrates. Biochemistry 38, 3837–3849 (1999).

32. J. Gruszczyk et al., Cryo-EM structure of the agonist-bound Hsp90-XAP2-AHR cytosolic complex. Nat Commun 13, 7010 (2022).

33. S. García-Alonso et al., Structure of the RAF1-HSP90-CDC37 complex reveals the basis of RAF1 regulation. Mol Cell 82, 3438–3452.e8 (2022).

34. A. K. Shiau, S. F. Harris, D. R. Southworth, D. A. Agard, Structural Analysis of E. coli hsp90 reveals dramatic nucleotide-dependent conformational rearrangements. Cell 127, 329–340 (2006).

35. I. Drulyte et al., Approaches to altering particle distributions in cryo-electron microscopy sample preparation. Acta Crystallogr D Struct Biol 74, 560–571 (2018).

36. K. A. Verba et al., Atomic structure of Hsp90-Cdc37-Cdk4 reveals that Hsp90 traps and stabilizes an unfolded kinase. Science 352, 1542–1547 (2016).

37. F. C. Fiesel, E. D. James, R. Hudec, W. Springer, Mitochondrial targeted HSP90 inhibitor Gamitrinib-TPP (G-TPP) induces PINK1/Parkin-dependent mitophagy. Oncotarget 8, 106233–106248 (2017).

38. R. Gómez-Sánchez et al., Mitochondrial impairment increases FL-PINK1 levels by calcium-dependent gene expression. Neurobiol Dis 62, 426–440 (2014).

39. C. Zhang et al., The plant triterpenoid celastrol blocks PINK1-dependent mitophagy by disrupting PINK1’s association with the mitochondrial protein TOM20. J Biol Chem 294, 7472–7487 (2019).

40. D. M. Bjorklund et al., Recognition of BRAF by CDC37 and Re-Evaluation of the Activation Mechanism for the Class 2 BRAF-L597R Mutant. Biomolecules 12, 905 (2022).

41. J. Doostzadeh, J. W. Tetrud, M. Allen-Auerbach, J. W. Langston, B. Schüle, Novel features in a patient homozygous for the L347P mutation in the PINK1 gene. Parkinsonism Relat Disord 13, 359–361 (2007).

42. Y. Hatano et al., Novel PINK1 mutations in early-onset parkinsonism. Ann Neurol 56, 424–427 (2004).

43. J. Shao, T. Prince, S. D. Hartson, R. L. Matts, Phosphorylation of serine 13 is required for the proper function of the Hsp90 co-chaperone, Cdc37. J Biol Chem 278, 38117–38120 (2003).

44. C. K. Vaughan et al., Hsp90-dependent activation of protein kinases is regulated by chaperone-targeted dephosphorylation of Cdc37. Mol Cell 31, 886–895 (2008).

45. M. Jaime-Garza et al., Hsp90 provides a platform for kinase dephosphorylation by PP5. Nat Commun 14, 2197 (2023).

46. B. Panaretou et al., Activation of the ATPase activity of hsp90 by the stress-regulated cochaperone aha1. Mol Cell 10, 1307–1318 (2002).

47. Y. Liu et al., Cryo-EM structures reveal a multistep mechanism of Hsp90 activation by co-chaperone Aha1. bioRxiv 10.1101/2020.06.30.180695 (2020).

48. A. Punjani, J. L. Rubinstein, D. J. Fleet, M. A. Brubaker, cryoSPARC: algorithms for rapid unsupervised cryo-EM structure determination. Nat Methods 14, 290–296 (2017).

49. T. Bepler et al., Positive-unlabeled convolutional neural networks for particle picking in cryo- electron micrographs. Nat Methods 16, 1153–1160 (2019).

50. A. Punjani, D. J. Fleet, 3D variability analysis: Resolving continuous flexibility and discrete heterogeneity from single particle cryo-EM. J Struct Biol 213, 107702 (2021).

51. P. Emsley, K. Cowtan, Coot: model-building tools for molecular graphics. Acta Crystallogr D Biol Crystallogr 60, 2126–2132 (2004).

52. E. C. Meng et al., UCSF ChimeraX: Tools for structure building and analysis. Protein Sci 32, e4792 (2023).

53. P. D. Adams et al., PHENIX: a comprehensive Python-based system for macromolecular structure solution. Acta Crystallogr D Biol Crystallogr 66, 213–221 (2010).

