## Supplementary Information for "Structural basis for the folding of PINK1 by the HSP90–CDC37 chaperone complex"

**Supplementary Figure 1. Purification of the intact human PINK1–HSP90β–CDC37 complex.**

**(a)** SDS-PAGE analysis of the eluate from Ni-NTA affinity purification. **(b)** Gel-filtration chromatography and SDS-PAGE analysis. The fractions corresponding to the region marked by the red arrow in the chromatography profile were subjected to SDS-PAGE. The arrowhead in the profile and the black dashed box in the gel image indicate the fraction used for cryo-EM analysis.

**Supplementary Figure 2. Cryo-EM data processing of the intact human PINK1–HSP90β–CDC37 complex.**

Data were processed using CryoSPARC. The final 3D reconstruction was obtained at a resolution of 3.05 Å and 3.04 Å, based on the gold-standard Fourier shell correlation (FSC) criterion at 0.143.

**Supplementary Figure 3. Purification of the human PINK1–HSP90β–CDC37 complex with chemical crosslinking.**

**(a)** SDS–PAGE analysis of the affinity-purified complex samples with and without crosslinking by 0.025% glutaraldehyde. **(b)** Gel-filtration chromatography and SDS-PAGE analysis of the crosslinked sample. The fractions corresponding to the region marked by the red arrow in the chromatography profile were subjected to SDS–PAGE. The arrowhead in the profile and the black dashed box in the gel image indicate the fraction used for cryo-EM analysis.

**Supplementary Figure 4. Cryo-EM data processing of the human PINK1–HSP90β–CDC37 complex with chemical crosslinking.**

Data were processed using CryoSPARC. The final 3D reconstruction was obtained at a resolution of 3.08 Å, based on the gold-standard Fourier shell correlation (FSC) criterion at 0.143. The local resolution values are shown as a color-coded density map.

**Supplementary Figure 5. 3D VA analysis of the human PINK1–HSP90β–CDC37 complex with chemical crosslinking.**

3D Variability Analysis revealed six clusters showing distinct positions of the $FCL^{HSP90}$ relative to the C-lobe and CTE of PINK1. The red arrowheads in cluster 2, 4, and 6 indicate that the density of the $FCL^{HSP90}$ is in close contact with the C-lobe of PINK1. The black arrows in cluster 1, 3, and 5 indicate that the density of the $FCL^{HSP90}$ is detached from the C-lobe of PINK1.

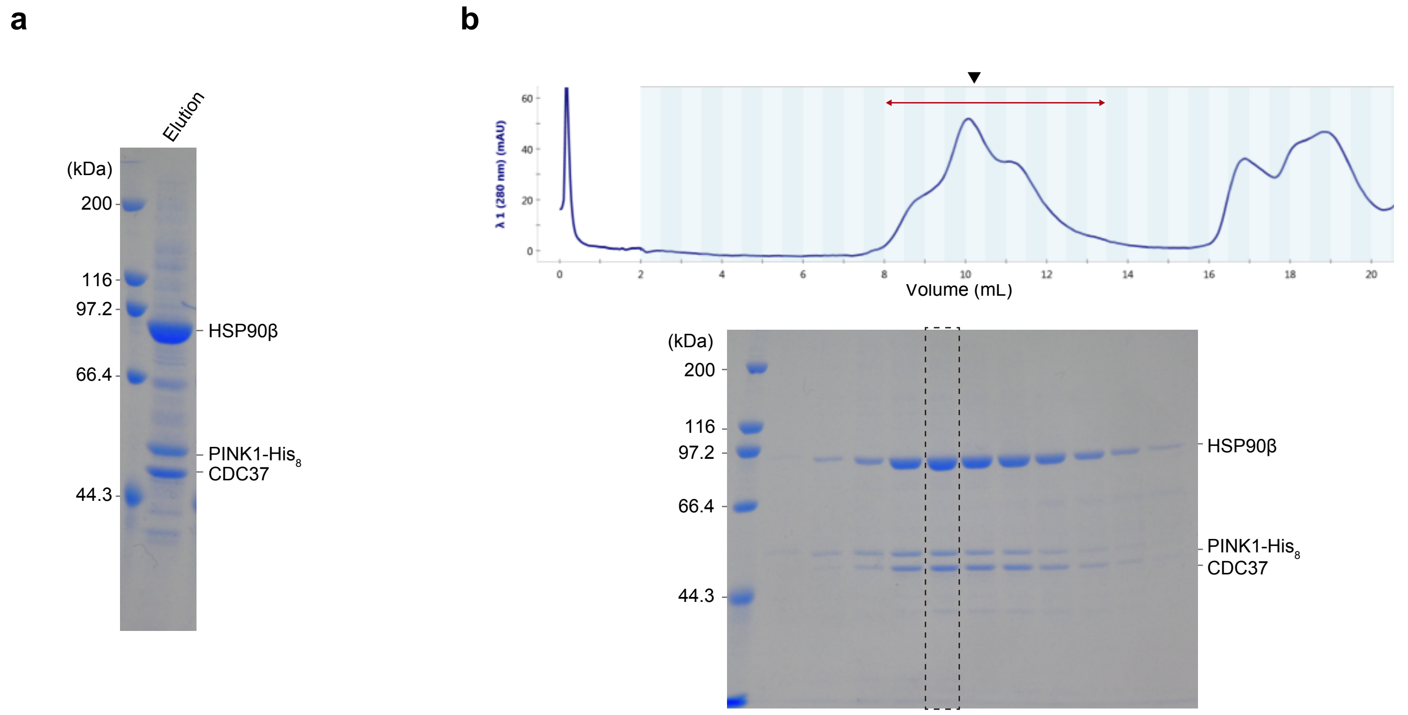


Okatsu et al. Supplementary Figure 1

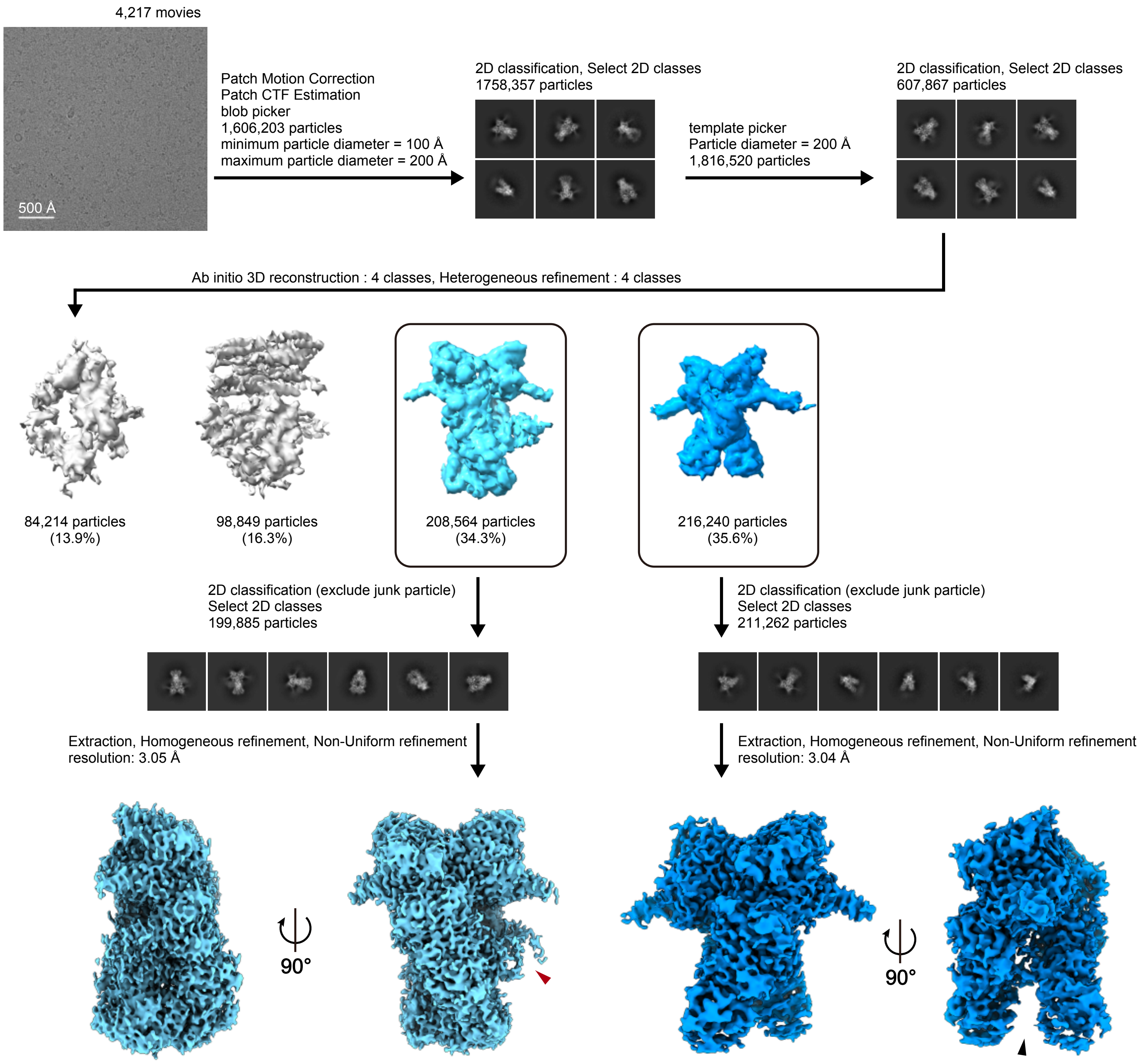


Okatsu et al. Supplementary Figure 2

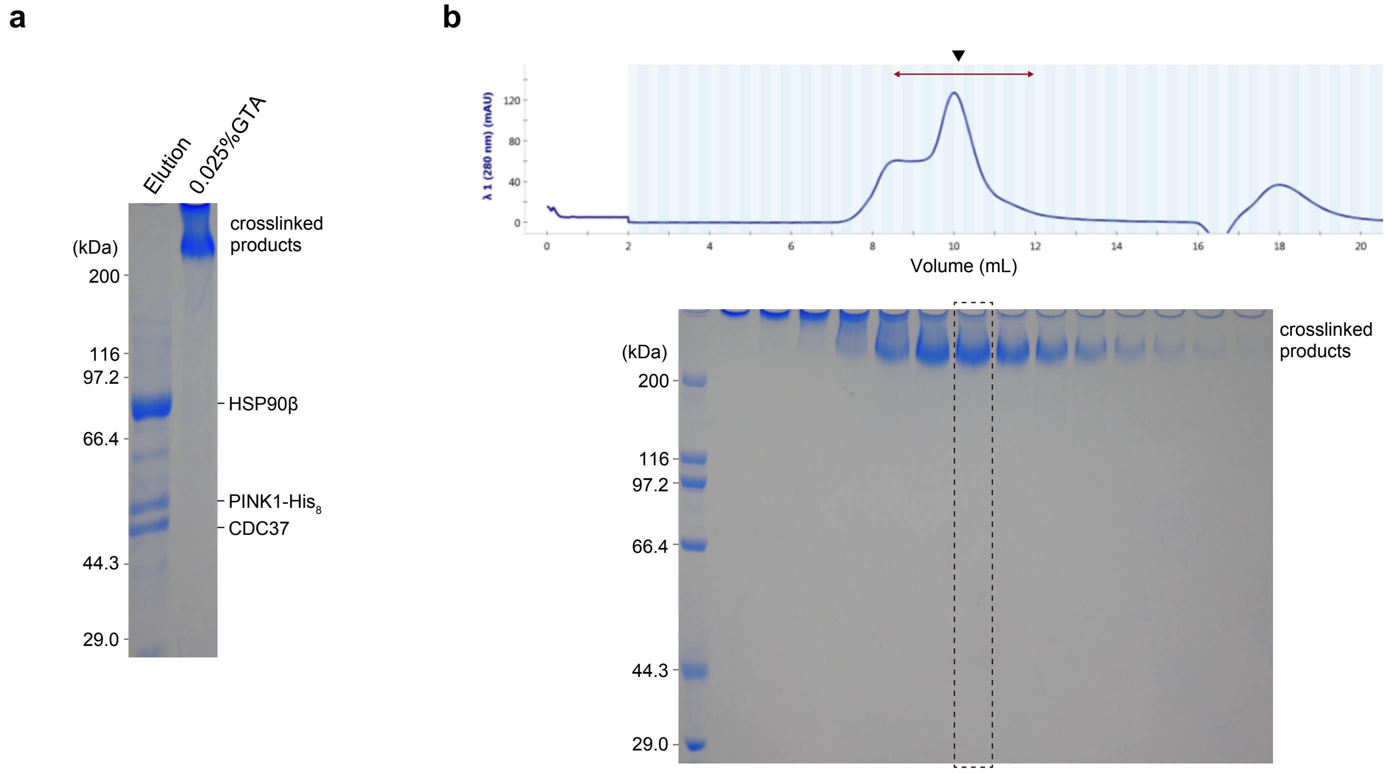


Okatsu et al. Supplementary Figure 3

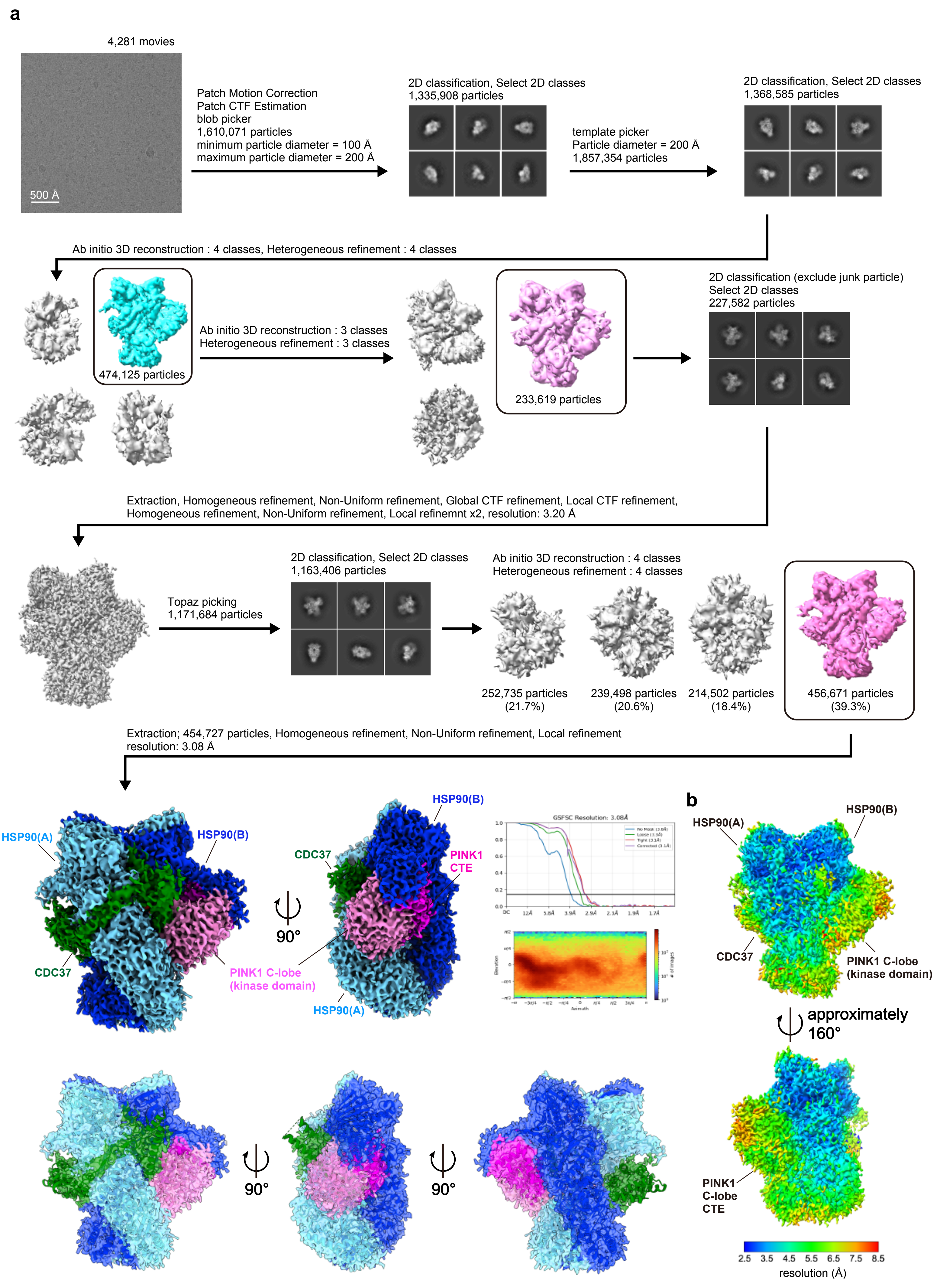


Okatsu et al. Supplementary Figure 4

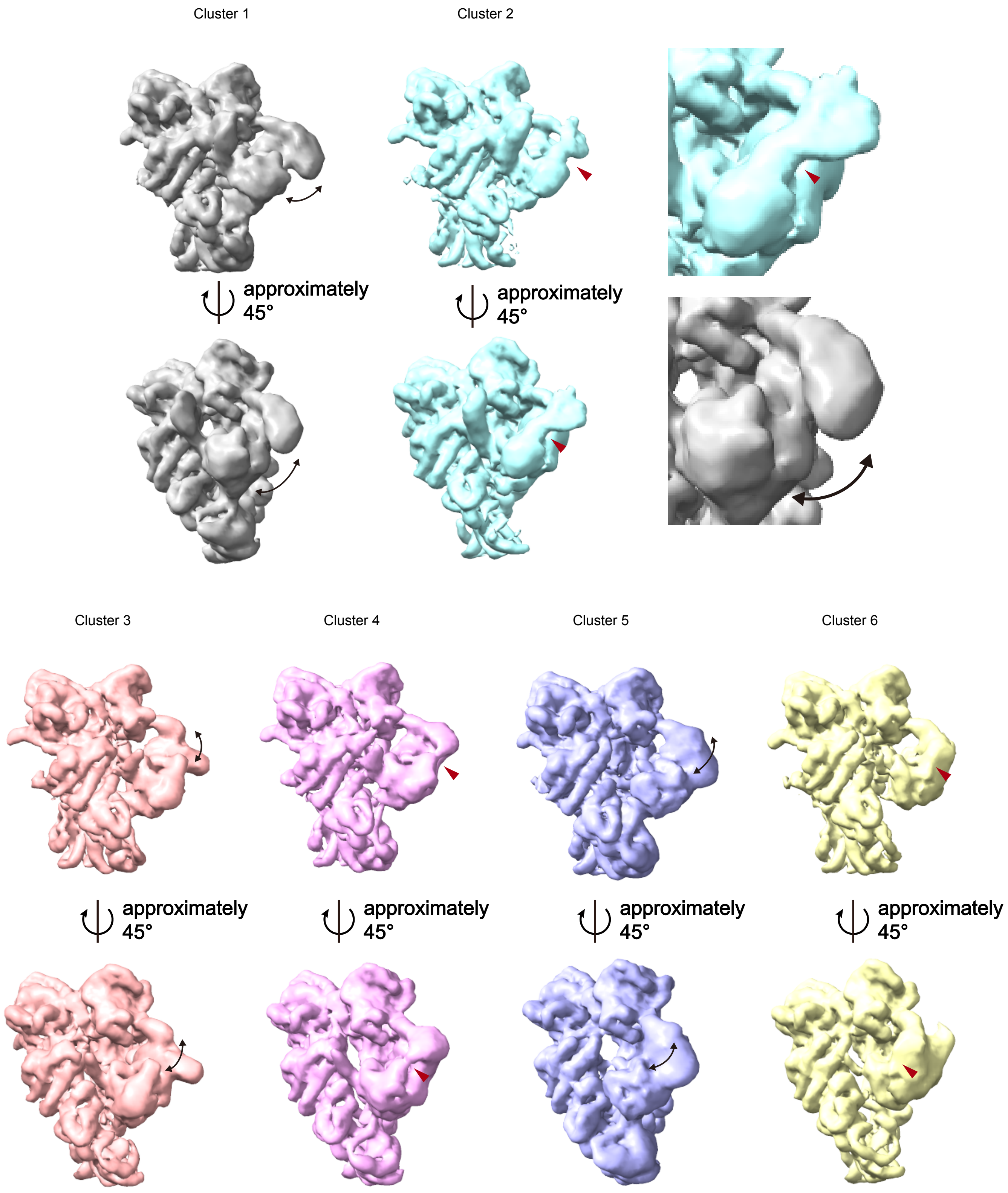


Okatsu et al. Supplementary Figure 5
